# Separation and enrichment of sodium-motile bacteria using cost-effective microfluidics

**DOI:** 10.1101/2020.11.24.395772

**Authors:** Jyoti P Gurung, Moein Navvab Kashani, Sanaz Agarwal, Gonzalo Peralta, Murat Gel, Matthew AB Baker

**Affiliations:** School of Biotechnology and Biomolecular Science, UNSW Sydney, NSW 2052, Australia; Future Industries Institute, University of South Australia, Mawson Lakes, SA 5095, Australia; Australian National Fabrication Facility – South Australia Node, Mawson Lakes, SA 5095, Australia; CSIRO Manufacturing, Clayton, South VIC 3169, Australia; CSIRO Future Science Platform for Synthetic Biology, Australia

**Author notes:** **Footnotes:** Author to whom correspondence should be addressed. Electronic mail.

## Abstract

Many motile bacteria are propelled by the rotation of flagellar filaments. This rotation is driven by a membrane protein known as the stator-complex, which drives the rotor of the bacterial flagellar motor. Torque generation is powered in most cases by proton transit through membrane protein complexes known as stators, with the next most common ionic power source being sodium. Sodium-powered stators can be studied through the use synthetic chimeric stators that combine parts of sodium- and proton-powered stator proteins. The most well studied example is the use of the sodium powered PomA-PotB chimeric stator unit in the naturally proton-powered *E. coli*. Here we designed a fluidics system at low cost for rapid prototyping to separate motile and non-motile populations of bacteria while varying the ionic composition of the media and thus the sodium-motive-force available to drive this chimeric flagellar motor. We measured separation efficiencies at varying ionic concentrations and confirmed using fluorescence that our device delivered eight-fold enrichment of the motile proportion of a mixed population. Furthermore, our results showed that we could select bacteria from reservoirs where sodium was not initially present. Overall, this technique can be used to implement selection of highly-motile fractions from mixed liquid cultures, with applications in directed evolution to investigate the adaptation of motility in bacterial ecosystems.

## I. INTRODUCTION

Halophilic, or salt-loving, microorganisms are predominantly found in salt-enriched habitats such as in the deep sea. One of the bacterial phenotypes that is directly influenced by the presence or absence of ionic salts such as sodium is bacterial motility^1^. Sodium-motility has particular medical relevance since pathogens such as *Vibrio cholerae*, of which there are up to 4M cases per year globally, strictly requires sodium ions to drive motility to spread disease^2^. Bacterial motility is, for the most part, driven by using a molecular motor called the bacterial flagellar motor (BFM). The BFM is a transmembrane nanomachine powered by cation influx consisting of a rotor, attached to a long filamentous protrusion known as a ‘flagellum’, and membrane-bound stator complexes that act as ion porters that couple ion transit to torque generation^3^. Bacteria navigate their surrounding medium by controlling when they change directions of rotating flagella, either clockwise or anticlockwise, to drive motility in a foraging search for nutrient, or to avoid repellents^4^.

The development of more efficient and cost-effective techniques to separate these bacteria with larger precision enables us to examine subtle differences in bacterial phenotype, in liquid and at high throughput. Microfluidics platforms are emerging as a technology with wide promise for microbiological research and development^5^ offering precise control of flow and buffer composition at the micron scales that bacteria inhabit. Microfluidic tools are in widespread use in combination with microscopy platforms and have been applied to the study and control of gradients which influence motility, to explore the mechanisms of chemotaxis, thermotaxis, rheotaxis, magnetotaxis and phototaxis^6^. Recent developments in microfluidics technology have enabled fine separation of cells based on subtle differences in motility traits and have applications in synthetic biology^7^, directed evolution^8^ and applied medical microbiology^9^.

In general, microfluidics platform consists of three essential components: (i) syringe pumps to generate flow inside the channel, (ii) a microfluidic chip containing micron-sized channels to direct flow, and (iii) observation and detection methods such as microscopy. Clean-room lithographic fabrication of channel moulds can be expensive, as are commercial microfluidic pumps. Recently cost-effective alternatives such as 3D printed microfluidic devices have been gaining more interest^10^. However, 3D printing of microfluidic channels with high spatial resolution and channel smoothness needs further development to completely replace other more standard microfabrication methods^11^. In addition, commercial pumps can be replaced by custom-built syringe pumps which are controlled by cheap micro-computers with strong open-source communities such as Arduino^12^. Such pumps perform well as demonstrated by a comparative study of commercial and Arduino-driven syringe pumps, which has shown no significant difference in the pump’s performance under measures such as stability of laminar flow^12^.

Most of the existing work in microfluidics for separating motile bacteria has been tested on proton-powered flagellar motors, and has predominantly focused on chemotaxis^6^. Here we wanted to explore the ability to build a simple fluidics device for screening mixed populations of bacteria for those motors which are powered from novel ionic power sources besides protons. Rather than testing the chemotactic response of the cell in response to attractant (culminating in phosphorylated CheY binding to the rotor to adjust switch frequency), we examined the capacity to separate motile, non-motile or partially motile cells based on an available energy source to power the stators to drive rotation. This would not be possible in native sodium-powered species such as *V. alginolyticus* as these cells cannot survive in low sodium, so we used our existing chimeric sodium-powered *E. coli* construct with which we have structural and functional familiarity^13^ to characterise the performance of our low-cost system.

We fabricated a simple and cost-effective microfluidics device with a single millimetre-sized channel with twin Y-shaped inlets and outlets. We demonstrated separation, selectionselection, and enrichment of sodium-motile bacteria (PomAPotB in *Escherichia coli*), using stable laminar-flow of two streams generated by pressure-driven flow. We characterised the performance of our system in varying sodium concentration and show the enrichment of the motile fraction of a mixed population and reconcile our measurements with simulation of active diffusion of sodium, motile, and non-motile bacteria.

## II. MATERIALS AND METHODS

### A. 3D printed custom-built syringe pump

Our custom-built syringe pump comprised of two 1 mL syringes in 3D printed syringe holders, two 28BYJ-48 (5V DC) stepper motors, an Arduino UNO R3 board, and an Adafruit Motorshield V2. Syringe adaptors were connected to the microfluidic chip using PTFE tubing (BOLA Inc.) with internal diameter (ID) of 0.5 mm and outer diameter (OD) of 1 mm. The stepper motors were connected to M1, M2, and M3, M4 ports on the Adafruit Motorshield V2 (Arduino UNO R3) and placed on the 3D printed syringe holder (Fig. 1c). A program based on C/C++ language was scripted to run two stepper motors at two independent rates simultaneously in the order of few hundreds of uL/min. The program code is presented in Supplementary information.

**Figure 1:**
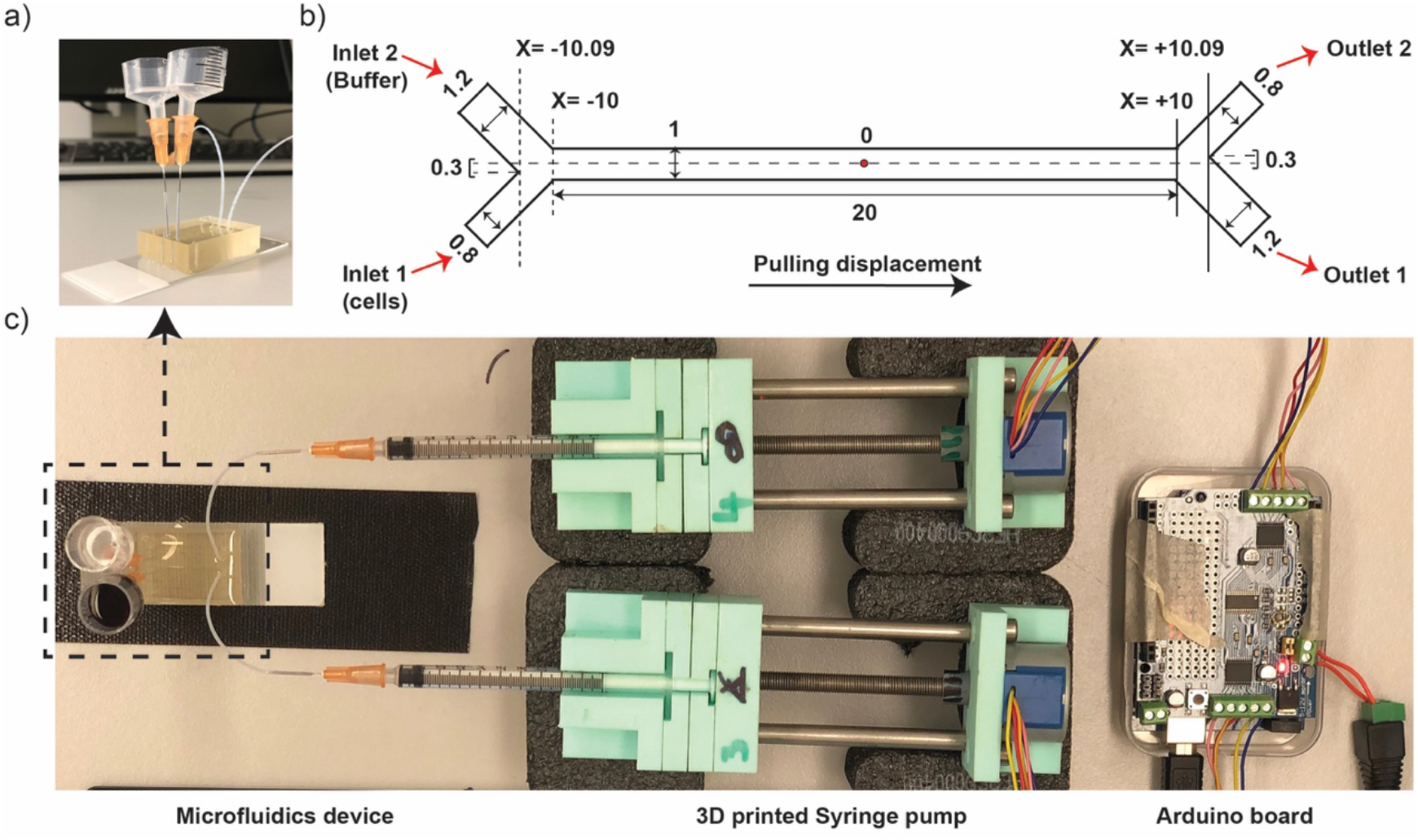
Fabrication of microfluidic device. a) Photo image of microfluidic device with upright reservoirs. b) Schematics diagram of asymmetric Y-shaped channel with unit of dimension in millimeters (mm) and origin point represented by ‘O’ letter in the center of the channel. c) Photo image of microfluidic platform containing Arduino board, custom-built 3D printed syringe pumps, and microfluidics device.

### B. Fabrication of microfluidics device

Y-shaped channel was designed using a CAD drawing tool, Fusion360 and the sketch was exported in .dxf file format to make it executable in a tape cutter (Silhouette Curio, USA). Brown adhesive tape was laminated on the glass slide (75 mm by 25 mm), baked in an oven at 65°C for 30 min, and cooled down to room temperature after gentle roll over the tape (removes air bubbles if present). The channel mould was cut on the slide using a tape cutter (Silhouette Curio, USA) and the excess tape outside of the channel structure was removed ^14^. Then, glass slide engraved with Y-shaped channel mould was placed in 3D-printed rectangular casket using tape (checked if the interfaces were tightly sealed well to prevent the leakage of PDMS), 10 g Clip-pack PDMS mix (SYLGARD®184) was poured into the casket and left in vacuum for degassing. PDMS was cured in oven at 65°C for 2 hours (or overnight). Cured PDMS was cut using scalpel and removed from the mould. Holes were punched at inlets and outlets using 0.75 mm biopsy punch (EMS/Rapid-core). Surfaces of PDMS cast and glass slide were treated with plasma discharge for 3 minutes and instantly bonded together with a slight press. Finally, microfluidics channel was treated with sterile 1% BSA solution (prepared in Milli-Q water) to prevent cell adhesion and stored at room temperature for future use.

### C. Image analysis of laminar-flow stability

Red dye was used to test the laminar flow stability inside the microfluidic channel. Initially, and at 30 min intervals during the microfluidics run, 5 minutes of video were recorded using Olympus inverted microscope at the magnification of 4X objective lens using ToupView imaging software. This 5 minute5-minute video was rendered to extract stack of image sequences using Photoshop (Adobe Inc.). Then, the image sequences were processed and analysed using Fiji (Image J) software to determine the laminar flow stability^12^. For this, a rectangular region of interest (ROI) was defined and cropped for a single image. Then, the contrast of cropped image was enhanced by 0.5%, black background was subtracted, converted to binary mask, and the area-fraction of red dye was measured. Ultimately, this operation for a single image was executed to a complete directory (other image sequences).

### D. Bacterial sample preparation

Plasmid pSHU1234 is based on pBAD33 backbone for expression of sodium-powered stator units; PomA and PotB whereas plasmid pSHOT is pSHU1234 but with complete deletion of PotB and partial deletion at the C-terminal of PomA from amino acid residue 203 to 254 (PomA1-D202-KLGCFGG-stop). These plasmids were transformed into *E. coli* strain RP6894; (ΔmotAmotB) to obtain a motile (pSHU) and non-motile (pSHOT) cell type respectively..respectively. These strains were subseqeuentlysubsequently co-transformed with plasmids, encoding fluorescent proteins for fluorescence experiments. RP6894 + pSHU1234 was transformed with DsRed.T4 plasmid^15^ (encodes red fluorescent protein); RP6894 + pSHOT was transformed with pACGFP1 plasmid (encodes green fluorescent protein, EGFP)^15^.

Approximately 4-5 colonies were picked from bacterial agar plates and inoculated into the LB broth. Then the culture was incubated in rotary shaker at 37°C for 2 hours (normally bacterial O.D. at 600 nm reaches the value > 0.5 after this time of incubation). Bacterial culture was centrifuged at 7000 rpm for 3 minutes, washed at least three times with water and finally suspended in motility buffer (10 mM KPi (pH 7.0), 85 mM NaCl, 0.1 mM EDTA-Na). Bacterial sample was resuspended in 1 mL volume to obtain the O.D. value between 0.5 – 0.6, and motility was checked under microscope before running inside microfluidics device.

### E. Operation of microfluidics device

500 uL of motility buffer or water was pushed into the channel to prefill the channel (Inlet 1 and Inlet 2). The device was then degassed in vacuum for around two hours to remove air bubbles. Before loading the sample, 300 uL of the solution was pulled out of the device, resulting in the inlet reservoirs being emptied, verifying smooth flow without clogging and leaving the channel prefilled. In first set of experiment, oOne of the reservoirs was loaded with red dye and the other with distilled water to characterise the stability of laminar flow in our microfluidic device. In second set of experiment, or either motile or non-motile bacterial sample suspended in motility buffer containing 85 mM of NaCl was loaded in one reservoir and the other with solution such as motility buffer containing 85 mM of NaCl. Finally in third set of experiment, to study the effect of Na+ ion diffusion in selecting sodium-motile bacteria, motile bacterial sample (i.e., pSHU bacteria) suspended in distilled water was loaded in one reservoir and other reservoir with (of varying concentration of NaCl (42.5 mM and 85 mM of NaCl)) or distilled water. Bacteria were collected from respective outlets, serially diluted, and 10 µL droplets were placed in the agar plate for colony count^16^. Here, separated fraction of motile bacteria was calculated as a ratio of bacterial count from Outlet 2 and total bacterial count from Outlet 2 and Outlet 2^17^. To demonstrate that the separated bacteria are viable and motile, the sample from Outlet 2 was spread plated for different dilutions in LB agar plates, replica plated into sodium-containing swim plates, and incubated at 37°C for 24 hours.

### F. Numerical simulation

Computational Fluid Dynamics (CFD) simulations were conducted to verify the performance of the microfluidic system and demonstrate the active diffusion of sodium, motile and non-motile bacteria in the device. In the simulations, the mediums were assumed to be an isothermal liquid, viscous, incompressible, and Newtonian fluids. Physical properties of the mediums are constant and unaffected by the channel geometry. The flow regime is laminar, and the gravity effects on the flow field are negligible.

Figure 1(b) shows the geometry used in this simulation and the Cartesian coordinate system that was placed in the center of channel length and width (red dot, Fig 1b). A uniform inlet velocity was set at the inlet boundary condition, and flow rate weighting of 1:2 (Outlet-2/Outlet-1) was placed at the outlet boundary condition. A no-slip boundary condition, where the velocity is zero, was applied at the walls in the computational geometry. The multi-zone quadrilateral-triangle computational mesh was generated in this domain using the pre-processor ANSYS Meshing software. The maximum mesh size used was 5 µm, and it is chosen based on a mesh independency analysis. No significant difference was observed in the simulations when the mesh was refined.

The computational platform used for the simulations was Fluid Flow (FLUENT) package of ANSYS (ANSYS Inc., 2019). For the isothermal flow of a Newtonian fluid of constant density, the governing equations were developed as incompressible Navier–StokesStokes’s equations, including momentum equation, and continuity equation. To include the species transport based on the concertation gradient in the diffusion-dominated laminar flow, the Maxwell-Stefan equations were used to find the diffusive mass fraction. The high degree Pressure-Implicit with Splitting of Operators (PISO) was used for pressure velocity coupling, and the PRESTO (Pressure Staggering Option) algorithm was employed for the pressure interpolation. The second-order upwind differencing and the Least Square Cell-Based gradient method was used for solving transport equations and the spatial discretization, respectively. The transient flow condition with bounded second-order implicit formulation was considered for simulation to monitor the flow within the device over time until it gets to the steady-state condition. The time step for the integration used is 10^−6^ second. In each time step, a maximum of 20 iterations was found to be sufficient for convergence, and the absolute convergence criterion was set to 10^−4^. The numerical simulations were carried out on a DELL Precision Tower having two processors of “Intel Xeon CPU E5-2699 V4”, including 25 physical cores with the maximum speed of 2.6 GHz for an individual core and 256 GB memory.

## III. RESULTS

### A. Fabrication and characterization of microfluidic device

A microfluidics device with Y-shaped inlets and outlets, connected to the main channel (length of 20 mm, width of 1 mm, and height of 0.065 mm) was fabricated to generate laminar flow and driven by Arduino-based 3D-printed syringe pump (Fig. 1c). The width of Y-shaped inlet and outlet were asymmetric, thus, one of the channel widths was 1.2 mm and other was 0.8 mm (Fig. 1b). This resulted the offset of 0.3 mm across the Y-axis (or channel width) about the line drawn along the channel length with origin in the mid-point of the main channel (Fig. 1b). Pulling displacement or negative pressure was applied using syringe pump at desired flow rates (Fig. 1b and c). We controlled flow rate by directing the interval time in milliseconds between steps of our stepper motor. Using 1 mL syringes, our Arduino system had a maximum flow rate of 50 µL/min, corresponding to driving the stepper motor at the maximum speed with 7.6 milliseconds interval between steps (Fig S1). Asymmetric geometry^15^ and unequal flow rates^17^ have been found to reduce the passing of bacteria across the laminar flow interface, and avoid collection of non-motile cells in Outlet 2. Thus, inlets and outlets of the device were arranged with asymmetric geometry in channel width (Fig. 1b) and we established a relative flow rate of 2:1 pulling through Outlet 1: Outlet 2. This ratio could be adjusted for the desired selection stringency. We measured the total volume flowed in a set amount of time (1 mL in 2 hours and 15 min in Outlet 1) and confirmed that a mean flow rate of 7.4 µL/min in Outlet 1 and 3.7 µL/min in Outlet 2 was sustained. Flow rates at inlet were equal, confirmed by equal decrement of inlet reservoir’s height throughout the experiment, and thus the flow rate inside the channel was assumed to be the average of 7.4 and 3.7 µL/min i.e., 5.55 µL/min across each stream (these flow rates for inlets and outlets along with the dimensions of the microfluidic channel were used for subsequent numerical simulation).

We characterized the system and stability of laminar flow using red dye to further implement for bacterial separation. Image analysis of 5 minute video at 30 minute intervals was recorded near inlet and outlet respectively to measure the stability of the laminar flow interface using red dye (Fig. 2). Fluctuation of red dye was measured as relative distance traversed by red dye away from the interface and was observed to be less than 10% of channel width i.e., ±0.05 mm for both inlets and outlets (Fig. 2).

**Figure 2:**
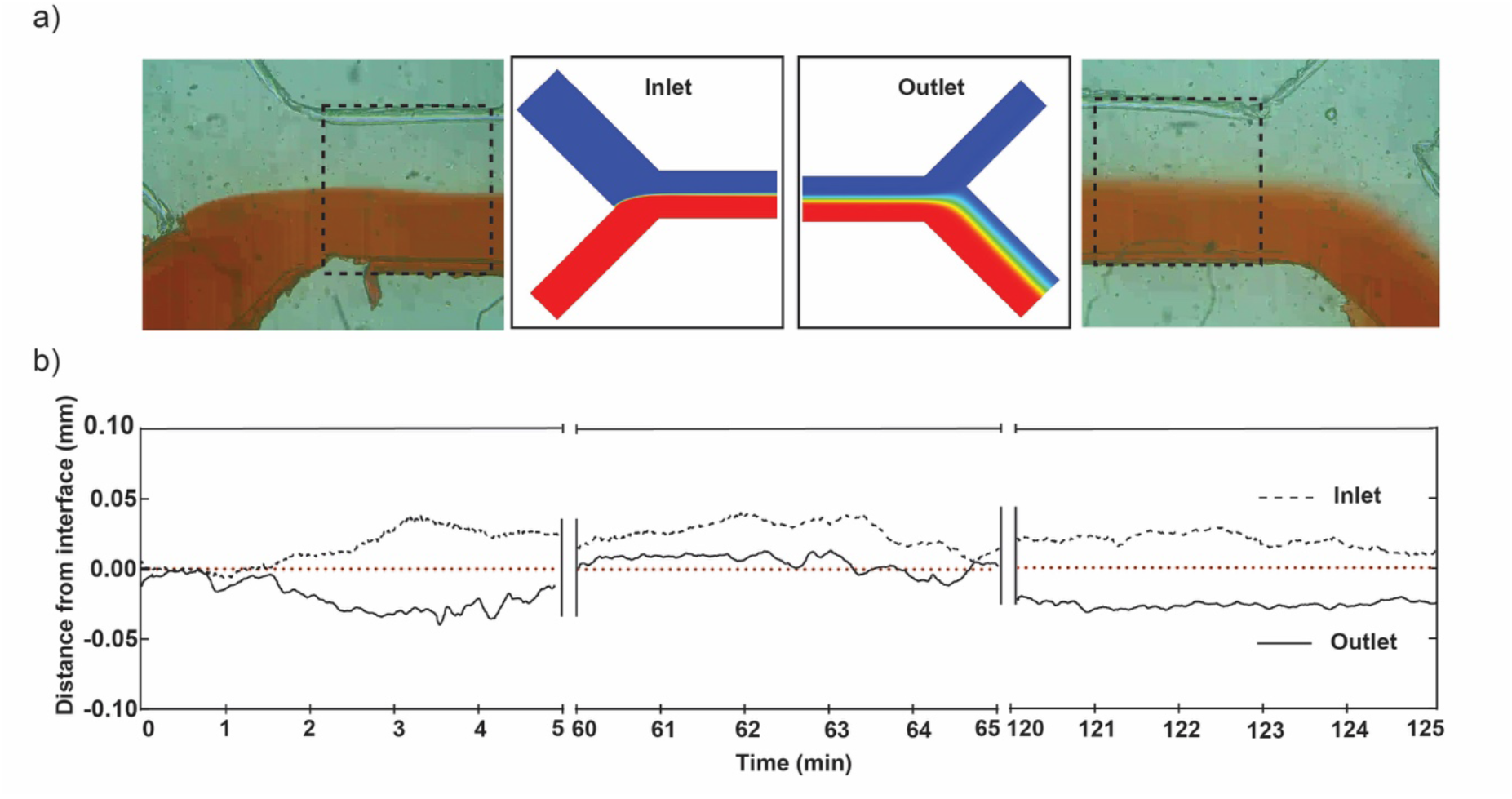
Characterization of microfluidic device. a) Laminar flow generated using our microfluidic device; represented by red dye (bottom) and colorless solution (water, top) streams in inlet (left) and outlet (right). Adjacent is numerical simulations for ion diffusion matched at inlet and outlet (see Fig 4). b) Line graph showing laminar interface fluctuation from center (0.00) over five-minute intervals to measure flow stability at 0 min, 60 min, and 120 min (plot of relative distance from interface vs time in minutes).

### B. Separation, selection and enrichment of sodium-motile bacteria

#### 1. Separation

We transformed a stator deleted *E. coli* strain (RP6894 ΔmotAmotB) with two different plasmids to prepare motile and non-motile bacteria. The sodium-motile strain was transformed with the pSHU plasmid (pSHU1234, chloramphenicol resistance, arabinose inducible pomApotB) which showed swimming in the presence of sodium ions. Our non-motile control was transformed with the pSHOT plasmid (chloramphenicol resistance, arabinose inducible PomA1-202 only, see Methods) (SI Fig. 3d). We drove the system for a total run time of 2 hours and 15 minutes to collect 1 mL of solution from Outlet 1 and 0.5 mL from Outlet 2 (see Materials and Methods). After the completion of a fluidic run, the output volumes from both outlets were serially diluted and plated to count the number of viable colonies after an overnight incubation (see Materials and Methods section). The separation efficiency was then measured as the ratio of number of bacteria from Outlet 2 to the total number of bacteria from Outlet 1 and Outlet 2 (i.e., S_eff_ = N_o2 /_ (N_o1_ + N_o2_))^17^. We observed the separation efficiency of (5±3) × 10^−5^ for non-motile pSHOT bacteria and (10±5) × 10^−4^ for sodium-powered motile pSHU bacteria (unpaired *t*-test p < 0.05). Motile pSHU bacteria which was suspended in 85 mM of Na+ ions (Inlet 1) and run against 85 mM of Na+ ions (Inlet 2) showed 30 times more separation than the non-motile pSHOT bacteria (Fig. 3a), confirming the efficiency of our device for separating and distinguishing motile and non-motile *E. coli*. (Fig 3a).

**Figure 3:**
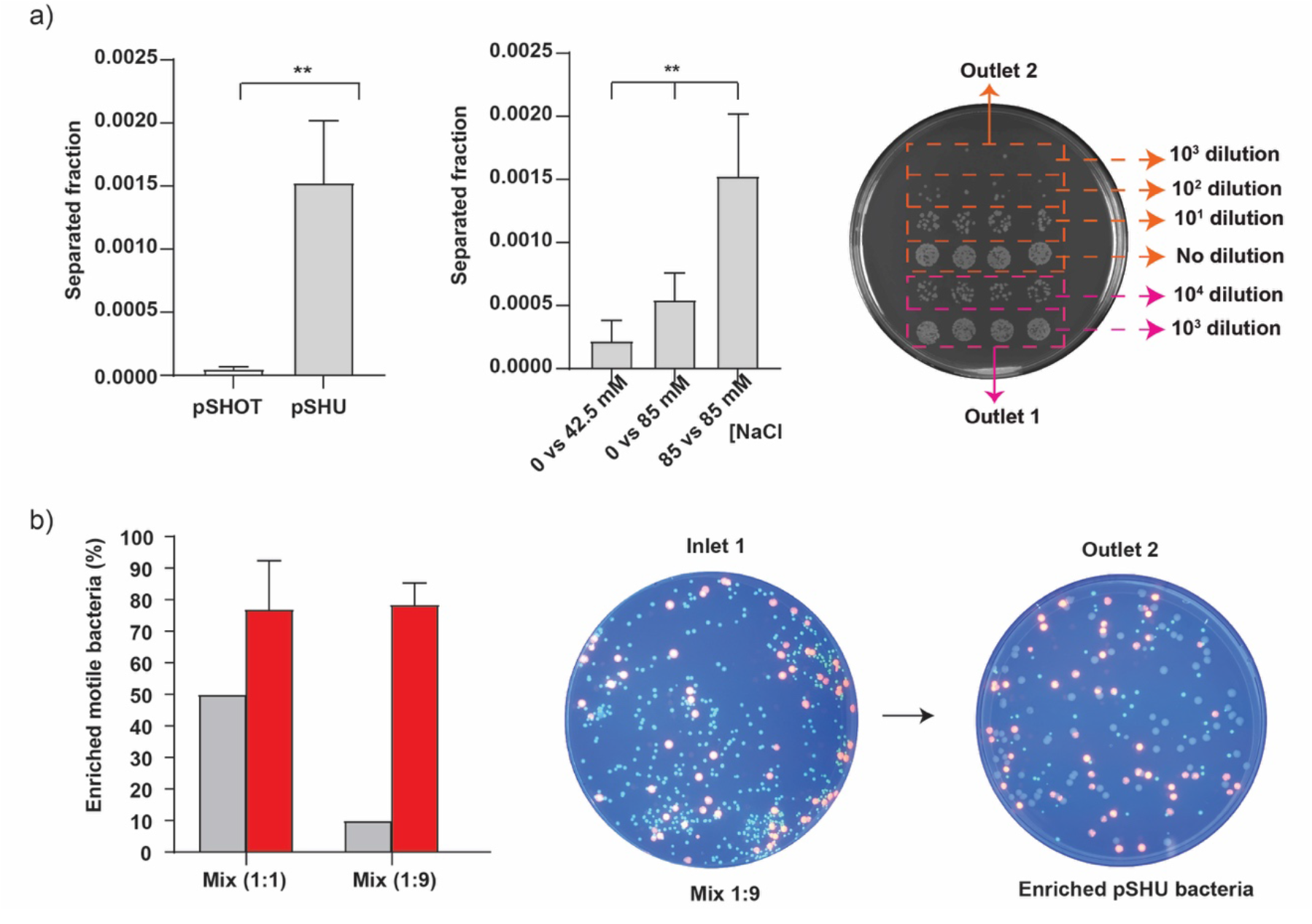
Separation and enrichment of sodium-motile bacteria. Inlet samples are configured as per Fig. 1. a) Left – separation of motile pSHU bacteria and non-motile pSHOT bacteria, suspended in motility buffer containing 85 mM of NaCl and run against the solution of motility buffer containing 85 mM NaCl, Middle – selection of motile pSHU bacteria, suspended in 0 mM of NaCl versus varying concentration of sodium (42.5 mM and 85 mM of NaCl), and Right – droplet method to count bacterial colony collected from Outlet 2 with no dilution, 10, and 10^2^ dilutions whereas Outlet 2 with 10^3^ and 10^4^ dilutions. b) Left - % of enriched motile pSHU bacteria for two different mixtures of fluorescent cells in inlets (pSHU:pSHOT; 1:1 and 1:9), Middle – Spread plate of fluorescent cell mix ration of 1:9 (10^4^ diluted) subjected into Inlet 1 of microfluidic device, and Right – Spread plate of 100 uL collected cells from Outlet 2 after separation (No dilution). Error bars represent standard deviation (SD) of triplicate data obtained from three independent experiments.

#### 2. Selection

We then used our device to test the capacity for selection of sodium-sensitive motile bacteria at low sodium concentration by varying the concentration of sodium ions. We loaded bacteria in buffer with no sodium ions (0 mM) (Inlet 1) and flowed in against 42.5 mM and 85 mM NaCl to test if diffusion of sodium from the high concentration flow stream (Inlet 2) would sufficiently drive rotation of flagella near the interface and enable separation and selection of motile pSHU bacteria. We observed increasing selection efficiency of motile pSHU bacteria with increasing sodium concentration: (3±2) × 10^−4^ for sodium ion concentration of 42.5 mM and (5±2) × 10^−4^ for 85 mM NaCl (Fig. 3a). Thus, the number of bacteria separated for 85 mM NaCl was approximately twice that for 42.5 mM NaCl, proportional to the external sodium concentration. Ordinary one-way ANOVA of selected pSHU bacteria in different concentration of Na^+^ ion showed separation efficiency was significantly different (p-value <0.05). As a control, we ran bacteria suspended in the buffer with no sodium at all in either of the inlets, which showed no separation for both non-motile pSHOT bacteria and motile pSHU bacteria (SI Fig. 3 – no viable colonies cultured from Outlet 2). The increasing selection efficiency of motile bacteria with increasing Na^+^ ion concentration is due to diffusion of sodium ions across the interface which was further supported by subsequent numerical simulation (Fig. 4).

**Figure 4:**
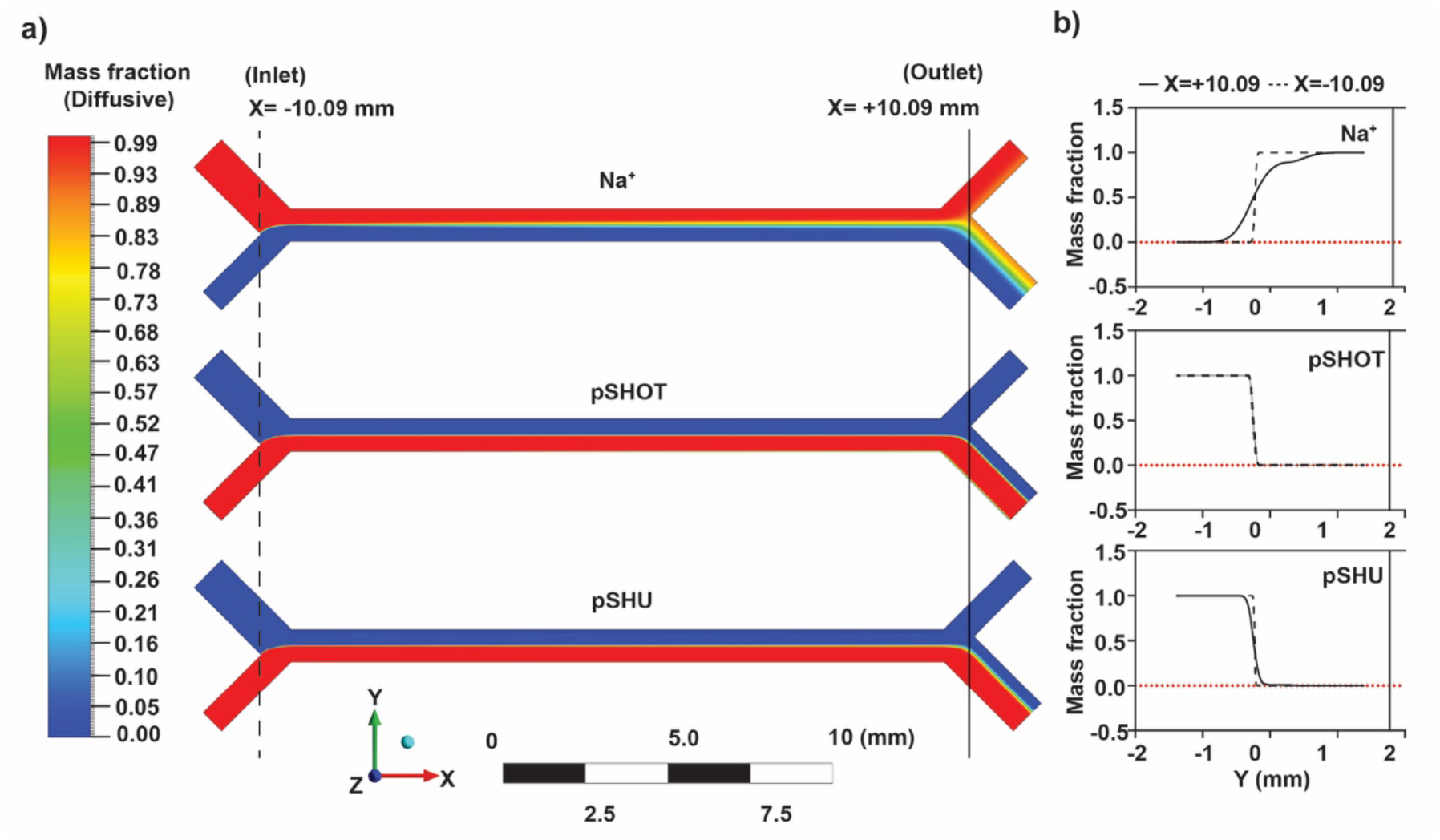
Numerical simulation using ANSYS Fluent software. a) (top to bottom) - image of simulation showing diffusion gradient of sodium ions, non-motile pSHOT, and motile pSHU. (Top to bottom) – Line graph of diffusive mass fraction vs distance in Y-axis (mm) for sodium ions, non-motile pSHOT, and motile-pSHU at the x-coordinate of ±10.09 mm from simulated data

#### 3. Enrichment

Finally, after separating and selecting motile bacteria, we used our microfluidic device to enrich sodium-motile bacteria from mixed populations of motile and non-motile bacteria. We further transformed RP6894 + pSHU with a plasmid for red fluorescent protein DsRed.T4 and RP6894 + pSHOT with a plasmid expressing green fluorescent protein EGFP (pACGFP1) (see Methods). To test our capacity to enrich a subpopulation and measure our enrichment efficiency, we prepared a mixed bacterial sample by mixing red fluorescent, motile cells and green fluorescent, non-motile, cells in the ratio of 1:1 and 1:9. We flowed these mixture solutions through our device as per our separation experiments above. The solution collected from Outlet 2 after fluidic run was then spread in agar plates to count and measure the enriched ratio of red and green colonies. In spread agar-plate, we observed three types of colonies in term of size and color i.e., large-sized red, small-sized green, and large-sized non-red colonies (SI Fig. 3). We confirmed that those non-red large-sized colonies are motile pSHU bacteria through sub-culturing on swim plates (SI Fig. 3). We observed that the percentage of separated sodium-motile pSHU bacteria was enriched from 50% to 77 ±16 % for 1:1 cell mix, and for the 1:9 cell mix, it was enriched from 10% to 79±7% (Fig. 3b). Thus, separation of motile bacteria for the initial cell-mixture ratio of 1:1 and 1:9 was approximately increased by 27% and 68% respectively (Fig 3b).

### C. Numerical simulation

We performed numerical simulations of our device to examine the diffusive spread of Na^+^ ions, motile, and non-motile cells. We approximated bacterial swimming as active diffusion, with diffusion coefficients of 8.3 × 10^−10^ m^2^/s (Na^+^), 5.32 × 10^−11^ m^2^/s (pSHU), and 2 × 10^−12^ m^2^/s (pSHOT) for Na^+^ ion^18^, motile bacteria, and non-motile bacteria^19^ respectively. The origin of the coordinate-space was defined as the midpoint of channel length and width (Fig 1b) and diffusive mass fraction was measured over the entire channel width (Fig 4). This mass fraction was further used to assess the profile across the channel at the inlet and outlet (at the distance of x = ±10.09 mm) to determine the separation at the divergence point, that is, the theoretical separation efficiency between Outlet 1 and Outlet 2.

Our simulation showed that Na^+^ ions diffused further across the interface than all types of bacteria during operation (Fig. 4), as expected from comparative size and diffusion input parameters. In our experiments we flowed 85 mM sodium ion into Inlet 1 (Fig 4A, top), and from the simulation of our flow conditions we calculated the sodium concentration at the midpoint of the channel (x = 0.00 mm, y = 0.00 mm) to be 39 mM. Thus, ∼3 mm across the interface, in the center of the channel, roughly half the sodium concentration is available to energize stators in cells to drive active diffusion.

Comparative bacterial diffusivity, proportional to the relative diffusion coefficients, was observed for motile pSHU and non-motile pSHOT bacteria (Fig 4b). The separation efficiency for the simulation was calculated using the ratio of mass fraction at the outlet point (x = +10.09 mm, y = + 0.03 mm). The separation efficiency of motile pSHU cells calculated from the simulation was 28 times greater than the simulation separation efficiency for non-motile pSHOT bacteria, in agreement with the separation efficiency from our experimental data i.e., 30 times.

## IV. DISCUSSION

There exist many microfluidics device for separating and sorting motile bacteria with high precision, accuracy, and efficiency according to various taxes^6^. However, many of those devices are fabricated using expensive microfabrication techniques which require sophisticated clean room facilities. Here we developed a simple and cost-effective microfluidic device using sellotape and PDMS, and Arduino powered syringe pumps to generate stable laminar flow in simple Y-shaped channel comparable to the other Arduino-based 3D printed syringe pumps^12^. Low cost separation has been achieved before, but many of these devices are designed to achieve high separation efficiencies at low flow rates^17^. Our low-cost device, in contrast, focuses on achieving high selectivity at low separation efficiency in order to select the most motile fraction of a larger population for subsequent culturing and examination for directed evolution^20^. Thus, we pursued low separation at the lowest flow rates at which we can achieve stable laminar flow and tuned our separation efficiency accordingly using a 2:1 ratio between cell and buffer streams. This ratio can be adjusted to increase or decrease the separation efficiency, or selectivity, accordingly.

We used pulling displacement via syringe pump in combination with upright reservoirs that maintained hydrostatic pressure (Fig. 1). This resulted in greater stability of the interface at both inlet and outlet (SI Fig. 5). Fluidic flow maintained via syringe pump can suffer from pressure drop (pulsation) along the channel. To reduce this, inlets were arranged with upright reservoirs of solution to impose hydrostatic pressure which acted as a pressure feedback system to minimize pulsation or pressure drop inside the channel (Fig 1a)^21^. We operated our system at high flow rates, resulting with low separation efficiency of 0.1% in comparison to other literature^17^. Use of higher flow rates was to reduce residence time for bacteria and reduce the number of non-motile bacteria that cross-contaminate our Outlet 2 sample (through advection with flow).

The benefits from using a fluidic system to measure and separate for motility are that we can select in-line from a liquid culture. Furthermore, we can recirculate the collected bacteria through the device till the point where we obtain the bacteria with highest degree of motility. The tunability of the selection, in combination with the recirculation approach could be further improved for long-term high-throughput motility screening of bacteria for applications alongside more sophisticated fluidics devices for directed evolution^22^.

Motile bacteria under the influence of flow tend to drift away from the flow direction due to helical motion of bacterial flagellar motor^23^. Inside rectangular microchannels, motile bacteria have shown drift both in the bulk fluid (rheotaxis) and upon interaction with surfaces nearby the channel boundaries^24^. In the opposite case, bacteria may get separated due to advection of fluid flow rather than. However, in our experiments, advection with flow should not be problematic:, at the mean flow rate of ∼ 5.55 µL/min, we have a system with low Reynolds number (Re ∼ 0.17) and high Peclet number (Pe _(Na+)_ ∼ 1715, Pe _(motile or pSHU)_ ∼ 26,750, and Pe _(non-motile or pSHOT)_ ∼ 711,540) signifying that the separation of motile bacteria in our device is due to active diffusion rather than advection or convection^25^. The negligible effect of flow advection in bacterial separation was further confirmed by our experimental observation where we did not see any cells arrive at Outlet 2 for bacteria in the absence of sodium (SI Fig. 3).

Previous microfluidics approaches to separation have observed a higher separation efficiency of ∼12% at flow rates similar to ours (∼ 5.55 µL/min, residence time ∼14 s) with the use of partitioning walls^17^. In our case our low separation efficiency was both by necessity and by design; we used relatively high flow rates as this was needed to maintain stable laminar flow in our system, and we also drove the streams in a 2:1 ratio to create a more selective system with lower separation efficiency. Our system did not have additional microstructures such as partitioning walls, and only controlled flow rates at the outlets, yet was able to reproduce a comparable purity of motile bacteria at separation^17^. In addition, we also verified the viability and motility of separated bacteria through subsequent culturing and motility plates (SI Fig. 3b). Separation would undoubtedly improve at lower flow rates, or with stronger active diffusion via concentration gradients of chemoattractants^15^, but at flow rates lower than 3.7 µL/min we observed transient instabilities in flow which resulted in the diversion of the cell-containing stream directly into Outlet 2, compromising our results.

We have shown here that sodium-powered motile bacteria can be selectively separated from a bacterial population even when suspended in sodium-free media. Diffused sodium ions from adjacent flow stream were able to power BFM rotation and induce motility in bacteria to then select for these motile bacteria. It is thus possible to use a similar approach to select for diverse bacterial species from mixed or unknown samples to screen for those cells that require alternative ions for motility such as potassium^26^ or even calcium^27^. As the separation efficiency of the device appeared correlated with ion concentration, we can adjust ion composition and flow rate to select for novel bacteria powered by rare or unusual ions. We can also use this approach to alternate ion sources to screen for bet-hedging bacteria^28^ that may be optimized to dual-power^29^.*onidiensis* or *Bacillus subtilis*

To measure enrichment of a motile population fraction, we used bacterial strains transformed with plasmids encoding fluorescent proteins (DsRed.T4 in motile pSHU bacteria and EGFP in non-motile pSHOT bacteria). We avoided using bacteria tagged with fluorescent motor proteins as these have been shown to affect the swimming speed of bacteria^30^, and would not be expressed at high copy number for low resolution detection. Bacterial populations, even those generated from a single colony, can show heterogenous phenotypes, including motility^31^, and indeed we observed a fraction of cells with no fluorescence, yet which we confirmed were still motile (SI Fig 3). We hypothesize this is due to the long maturation time required to properly fold dsRed to become fluorescent^32^. This also confounded attempts to use direct fluorescence measurements such as fluorescence intensity, as the overall intensity of a liquid volume was dependent on the amount of dsRed in the cytoplasm of the cells, which did not necessarily accurately reflect the number of motile cells in a liquid volume. The differently tagged cells also have differential growth rates (SI Fig. 3). This could explain the difference in colony size between small-sized colonies of non-motile green pSHOT bacteria and larger colonies of motile red pSHU bacteria (SI Fig. 3). This visual phenotypic difference could be used to verify pSHU vs pSHOT colony assignment during colony counting to determine the enrichment ratios in our experiments.

Separation efficiencies measured from diffusional simulation data of motile pSHU and non-motile pSHOT bacteria showed that motile pSHU diffused to Outlet 2 at a rate 28 times greater than non-motile pSHOT. This simulated separated efficiency data agreed with our experimental data (i.e., 30 times more for pSHU compared to pSHOT). The small discrepancy could arise from a few small differences: we only detect viable cells, not all cells, and undoubtedly lose some cells to surface coating of the tubing, and containers used in dilution and subsequent culturing. It is also known that bacterial swimming deviates from strictly diffusive behavior^17^, and the flow in our device is likely further from ideal behaviour in regions close to the outlets. Simulations offer benefit though comparison between the diffusivity of the energy source (sodium ions), with attractants, and with the diffusive capacity of motile and non-motile cells. This provides a mechanism to refine designs and geometries for specific applications in bacterial selection and separation.

## V. CONCLUSIONS

Our system can be rapidly prototyped and can separate small differences in diffusivity by tweaking flow rates and channel geometry for optimization of specific applications in micron-scale separation. In this paper, we fabricated a cheap, effective microfluidics system for separation and enrichment of sodium-powered motile bacteria from non-motile populations. Our system has broad applications in environmental and field screening of bacterial species from liquid, as well as for quick assays to test the bulk motility of a sample. We envision future applications in directed and experimental evolution, particularly where these fields intersect with the evolution and adaptation of bacterial motility. That said, this device is equally translatable for applications in other systems where micron-scale motility is of importance, such as in the separation of motile eukaryotes (such as sperm for fertility applications, or cilia) and the multiple motile archaea strains (powered by the archaellum) that are increasingly of interest in studies in convergent evolution and for harvesting novel biotechnological products.

## AUTHORS CONTRIBUTION

JY designed and executed fluidics, cell culturing and fluorescence experiments, and wrote the manuscript. MNK executed simulations and wrote the manuscript. SA designed, wrote, and tested code to drive Arduino. GP prepared the pSHOT construct. MG designed experiments and wrote the manuscript. MABB supervised the project, designed, and executed experiments, and wrote the manuscript.

## CONFLICT OF INTEREST

There are no conflicts to declare.

## SUPPLEMENTARY INFORMATION

See Supplementary Information for Figures 1, 2, 3 and 4.

## ACKNOWLEDGEMENTS

We would like to acknowledge the gift of a 3D printed holder for the syringe pumps and a test Arduino from Dr James Walsh and A/Prof Till Boecking from the Centre for Single Molecule Science, UNSW Sydney. We also would like to thank Prof. Wenbin Du from the Institute of Microbiology at the Chinese Academy of Sciences, Beijing, for the plasmid encoding for fluorescent proteins used in this work. This work was conducted (in part) using the ‘Design House’ facilities at the South Australia node of the Australian National Fabrication Facility (ANFF-SA), a company established under the National Collaborative Research Infrastructure Strategy to provide nano- and micro-fabrication facilities for Australia’s researchers. MABB was supported by a UNSW Scientia Research Fellowship, a CSIRO Synthetic Biology Future Science Platform 2018 Project Grant, and ARC Discovery Project DP190100497.

## DATA STATEMENT

The data that supports the findings of this study are available within the article and its supplementary material.

## Notes

### Competing Interest Statement

The authors have declared no competing interest.

### Summary of Updates

Updated text following journal revisions. New Supplementary Figure 5.

