## Supplementary Figures 1-5 for "Separation and enrichment of sodium-motile bacteria using cost-effective microfluidics"

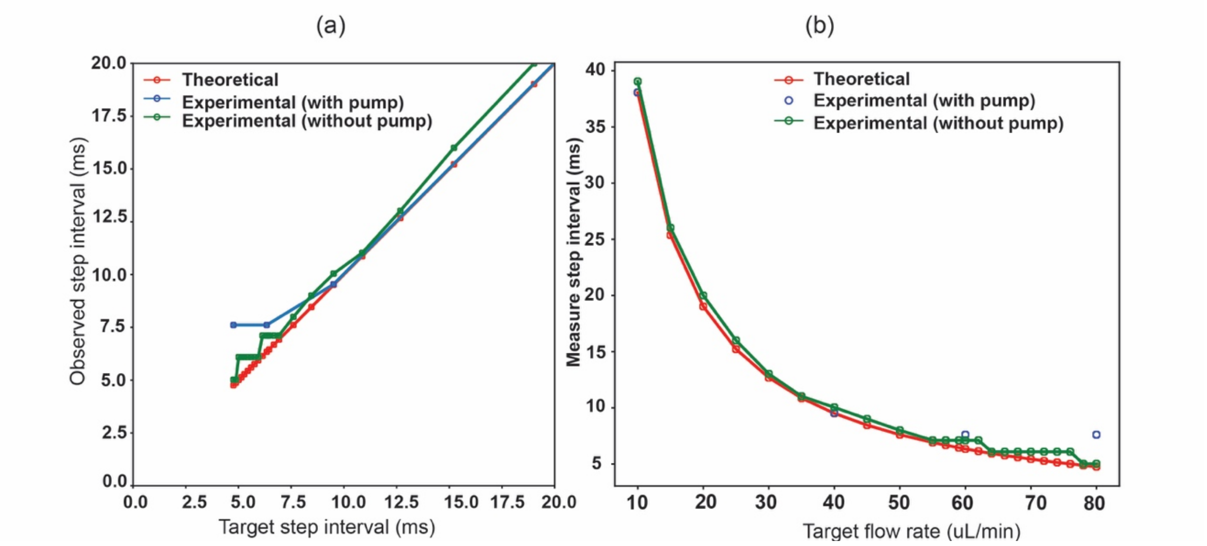


**SI Figure 1:** Step interval comparison of computation target value and experimentally measured value (both with and without syringe pump). a) Line graph of observed step interval (ms) versus target step interval (ms). b) Line graph of step interval (ms) vs flow rate (µL/min). Below 10 ms interval, corresponding to a flow rate of 35 uL /min the target and the observed values diverge.

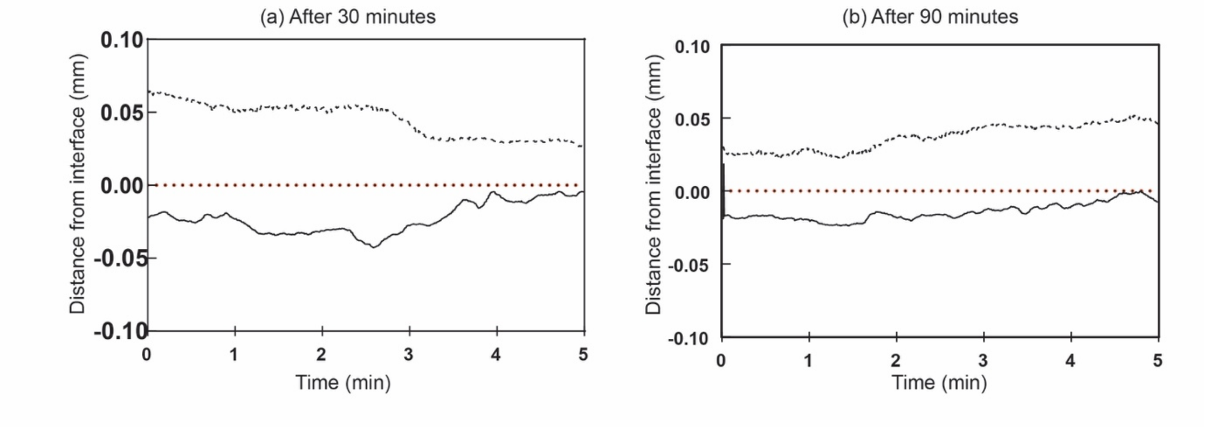


**SI Figure 2:** Line graph of relative interface from interface vs time (min). a) after 30 minutes of fluidic run. b) after 90 minutes of fluidic run.

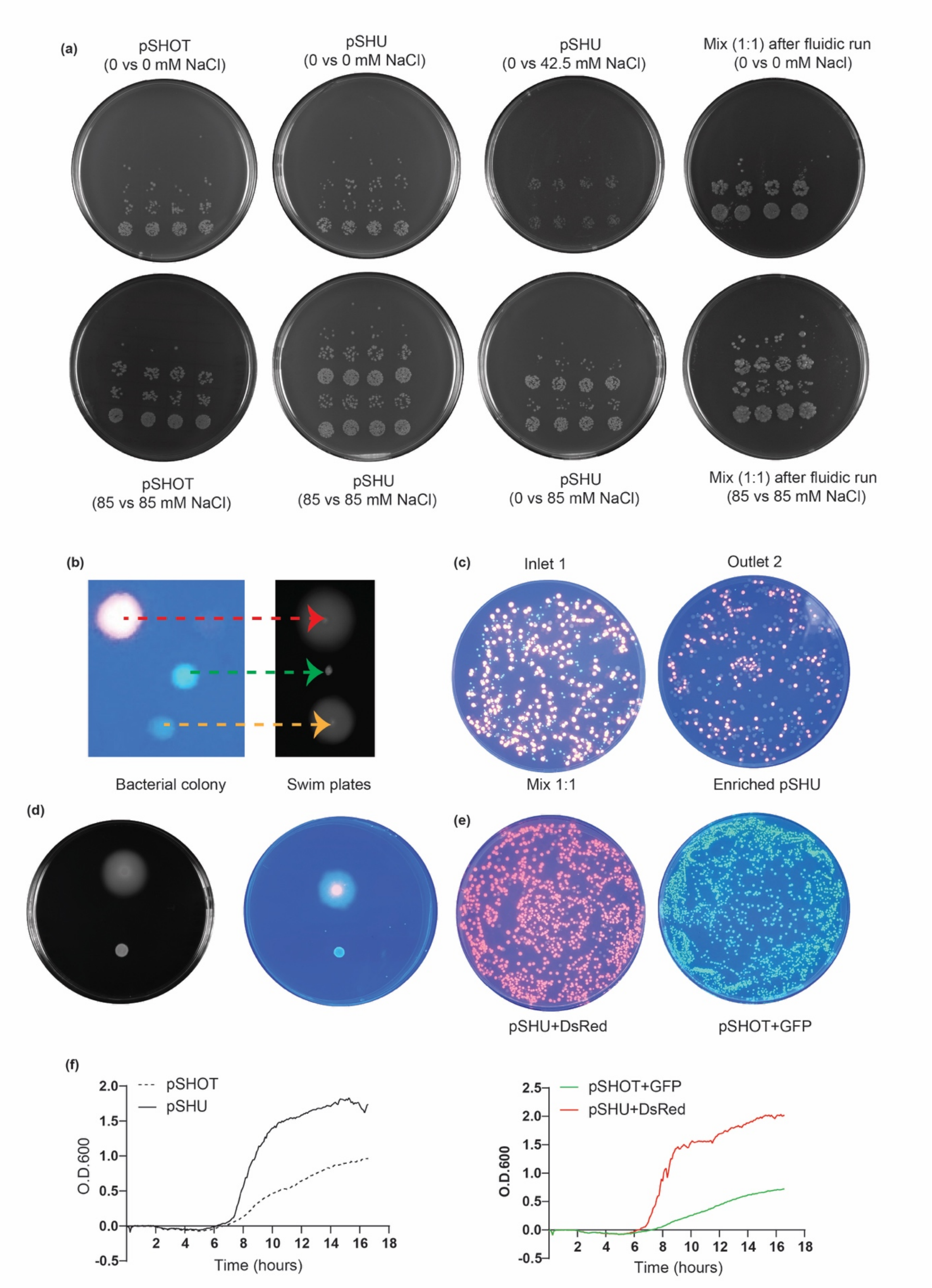


**SI Figure 3:** a) Images of agar plates used for bacterial colony count by droplet method at different sodium ion concentration. b) Swim plate of red, non-red, and green bacterial colonies. c) UV illuminated image of spread plates for mix 1:1 in inlet 1 ($10^4$ dilution) and outlet 2 (no dilutions). d) Left - swim plate of pSHU and pSHOT, Right – swim plate of pSHU + DsRed.T4 (red fluorescent) and pSHOT + EGFP (green, fluorescent). e) Spread plate of red fluorescent pSHU +DsRed.T4 and green, fluorescent pSHOT + EGFP. f) Growth curve of pSHOT and pSHU (left), pSHOT + EGFP and pSHU + DsRed.T4.

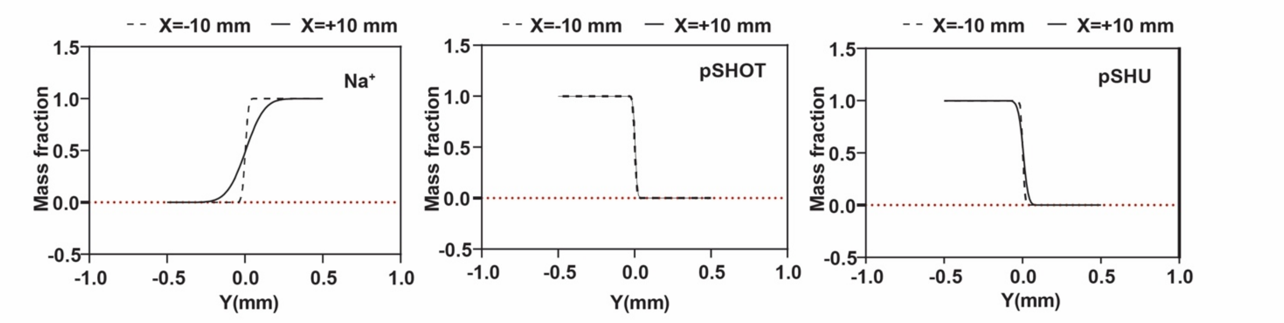


**SI Figure 4:** Line graph of diffusive mass fraction vs distance in Y-axis (mm) for sodium ions (left), pSHOT (middle), and pSHU (right) at the x-coordinate of distance ±10 mm.

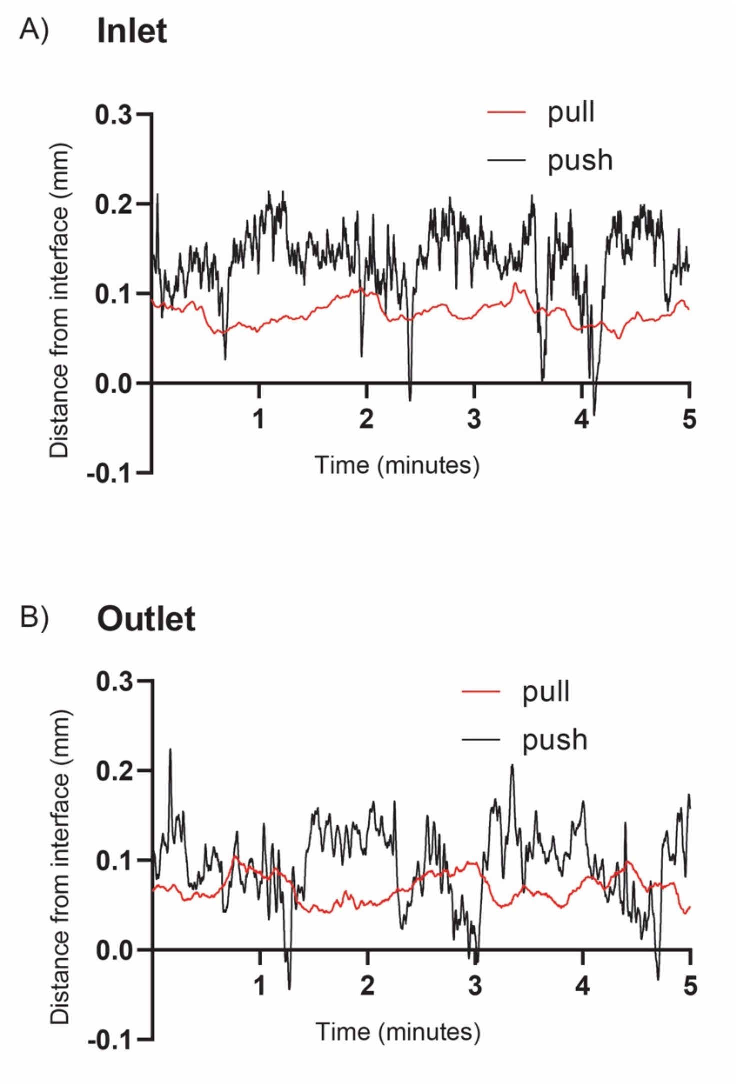


**SI Figure 5:** Line graph of relative interface from interface vs time (min) comparing pulling vs pushing driven fluidics. A) Distance from interface at inlet (0 minutes after starting flow) comparing push (black) and pull (red) displacement respectively. B) Distance from interface at outlet (0 minutes after starting flow) comparing push (black) and pull (red) displacement respectively.
